# Major cell-types in multiomic single-nucleus datasets impact statistical modeling of links between regulatory sequences and target genes

**DOI:** 10.1101/2022.09.15.507748

**Authors:** F. JA Leblanc, G. Lettre

## Abstract

Most variants identified by genome-wide association studies (GWAS) are located in non-coding regions of the genome. While largely untested functionally, it is assumed that most of these GWAS variants modulate the activity of enhancers. Epigenomic profiling, including ATACseq, is one of the main tools used to define enhancers. Because enhancers are overwhelmingly cell-type specific, inference of their activity is greatly limited in complex tissues that include multiple cell-types. Multiomic assays that probe in the same nucleus both the open chromatin landscape and gene expression levels enable the study of correlations (links) between these two modalities. Current best practices to infer the regulatory effect of candidate *cis*-regulatory elements (cCREs) in multiomic data involve removing biases associated with peak coverage and GC content by generating null distributions of matched ATACseq peaks drawn from different chromosomes. This is done under the assumption that the tested *cis*- and the matched *trans*-ATACseq peaks are uncorrelated. This strategy has been broadly adopted by popular single-nucleus multiomic workflows such as Signac. Here, we uncovered limitations and confounders of this approach. We found a strong loss of power to detect a regulatory effect for cCREs with high read counts in the dominant cell-type. We showed that this is largely due to cell-type-specific *trans*-ATACseq peak correlations creating bimodal null distributions. We tested alternative models and concluded that physical distance and/or the raw Pearson correlation coefficients are the best predictors for peak-gene links when compared to predictions from Epimap (e.g. CD14 area under the curve [AUC] = 0.51 with the method implemented in Signac vs 0.71 with the Pearson correlation coefficients) or validation by CRISPR perturbations (AUC = 0.63 vs 0.73).

## Introduction

Understanding how the non-coding genome regulates gene expression is paramount to attribute functions to noncoding variants identified by genome-wide association studies (GWAS). We can gain insights into the regulatory potential of non-coding regions through epigenetic mark assessments (CHIPseq), chromatin conformation capture methods (3C, Hi-C, ChIA-PET), expression quantitative loci analysis (eQTL), and open chromatin sequencing (ATACseq and DNase-seq). Extensive databases that collate and summarize these methods’ results in a broad range of cell lines and tissues are now available (i.e. ENCODE^1^, GENECARD^2^).

Leveraging on this data, statistical models such as those derived by the activity by contact^3^ (ABC) method and the Epimap^4^ project have generated strong predictions about the regulatory potential of many non-coding regions. Some of these predictions have been experimentally validated by CRISPR screens, often carried out in cancer cell lines^3,5^.

Direct measurement of both open chromatin regions and gene expression can be done concomitantly within a single nucleus using multiomic methods. At the single-nucleus resolution, the correlation of ATACseq peaks and RNAseq genes read counts (henceforth defined as links) provides highly specific hypotheses about the regulatory potential of non-coding candidate *cis*-regulatory elements (cCRE). Current best practices to analyze multiomic datasets and infer the regulatory effect of cCRE involve removing biases associated with ATACseq peak coverage and GC content. As proposed by Ma et al. 2020^6^, that method builds a null distribution of gene-peak correlations using ATACseq peaks of matching coverage and GC content drawn from chromosomes excluding the one hosting the tested gene (*trans*-links). The resulting distribution of Pearson correlation coefficients is then scaled, providing Z-scores for the *cis*- and each matched *trans*-links (**Fig. 1A**). This is done under the assumption that these *trans*-ATACseq peaks should not have a regulatory effect on the tested gene. This strategy has been broadly adopted by popular single-nucleus multiomic workflows such as Signac^7^.

**Figure 1.**
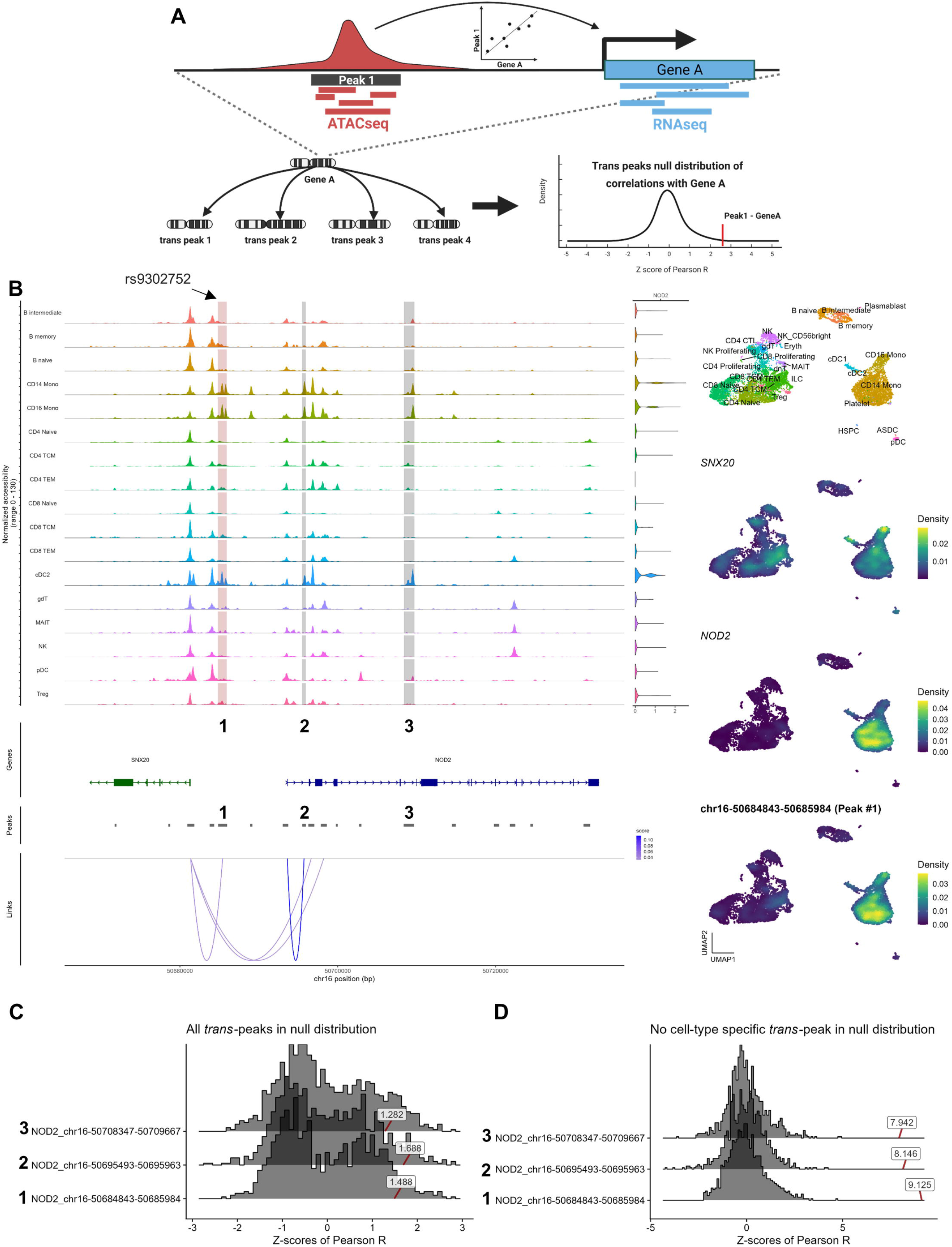
The Z-scores method misses candidate regulatory sequences linked to *NOD2* expression in peripheral blood mononuclear cells (PBMC). (**A**) The Z-scores model matches an ATACseq peak for GC content and coverage with ATACseq peaks in *trans* to create a scaled null distribution, producing Z-scores for each *trans*-links and the tested peak. (**B**) ATACseq tracks at the *NOD2* locus identified in PBMC. The grey areas (labeled 1-2-3) highlight the top three ATACseq peaks correlations with *NOD2* expression using the simple Pearson R method. Peak #1 (chr16-50684843-50685984) includes an eQTL for *NOD2* that is also associated with leprosy and Crohn’s disease by GWAS. The loops highlighted in the “Links” row are identified using the Z-score method (P-value <0.05); we note that there is no significant link between peak chr16-50684843-50685984 (peak #1) and *NOD2*. Loops are drawn from the middle of the ATACseq peaks to the transcription start site of the correlated gene(s). In the right column, we showed (top to bottom) RNAseq UMAP of cell-type annotations, *SNX20* expression density, *NOD2* expression density, and chr16-50684843-50685984 ATACseq accessibility density. The violin plots represent *NOD2* expression levels in each cell-type. (**C**) Three GC- and coverage-matched null distributions for ATACseq peaks (peaks #1-2-3) at the *NOD2* locus generated using the Z-scores method. Labeled boxes represent the corresponding Z-score statistics for the peaks tested against *NOD2* expression. Only peak #2 is significant using this approach (nominal P-value = 0.04). (**D**) As in **C**, but we generated the null distributions after excluding ATACseq peaks specific to the same cell-type as the tested ATACseq peak (see **Methods** for details). With this strategy, the three peaks (#1-2-3) are significantly linked with *NOD2* expression (P-value <1×10^−15^).

Here, by analyzing a publicly available multiomic peripheral blood mononuclear cells (PBMC) dataset (**Methods**), we uncovered limitations and confounders associated with this approach (termed the Z-scores method below). We found that the Z-scores method results in a strong loss of power to detect the regulatory effect of cCREs with high read counts in the most abundant cell-type(s). We tested various alternative models and concluded that the simplest approach, that is the raw Pearson correlation coefficients (this method is termed Pearson R below) and/or physical distance is computationally advantageous and provides the best predictions of “ATACseq peak-target gene” links when compared to results from Epimap or CRISPR perturbation screens.

## Results

### The number of cells in each cell-type biases the null distributions and statistics of the Z-scores method

In this study, we refer to Z-score as the scaled Pearson R value of a *cis*-link between an ATACseq peak and a nearby gene against its matched *trans*-link null distribution (the Z-scores method, **Fig. 1A**). After processing the PBMC multiomic data with Signac (**Fig. S1** and **Methods**), we noticed striking differences in terms of statistical significance for many peak-gene links when comparing the Pearson R coefficients and the Z-scores. For instance, the ATACseq peak chr16-50684843-50685984, upstream of *NOD2*, contains a *NOD2* eQTL (rs9302752) in whole blood, liver, tibial nerve, spleen and brain based on data from GTEx^8^ that is also associated with leprosy and Crohn’s disease by GWAS (**Fig. 1B)**^9^. In the PBMC dataset, this ATACseq peak and *NOD2* expression are relatively specific to monocytes and are correlated (R=0.12), although no significant links are identified using the Z-scores method (nominal P-value=0.07) (**Fig. 1B**). However, a significant link is identified between this peak and *SNX20*, a gene with an expression density that poorly overlaps the ATACseq signal at chr16-50684843-50685984 (**Fig. 1B**, right column). Importantly, we also noted that the null distributions for the *trans*-peaks matched with three ATACseq peaks at the *NOD2* locus are bimodal, making their corresponding peak-gene link Z-score statistics inaccurate (**Fig. 1C**). When we exclude from the dataset ATACseq peaks that are specific to the cell-type in which the ATACseq peak is mostly accessible and then create a null distribution with the remaining *trans*-peaks, the distribution is unimodal (**Fig. 1D**). This suggests that the choice of which ATACseq peaks are included in the null distribution has a huge impact on the Z-scores method results (see below).

Next, we performed several analyses to better understand how cell-type composition affect the identification of ATACseq peak-gene links from single-nucleus multiomic experiments. In the PBMC dataset, CD14 monocytes is the dominant cell-type (n=3075 cells [27%])(**Fig. 2A)** and clusters with CD16 monocytes and classical dendritic cells 2 (cDC2) both on ATACseq and RNAseq UMAP (**Fig. S1**). We refer to this cell archetype as mononuclear phagocytes (MP). We found that ATACseq peaks with specific accessibility in MP had lower median peak-gene link statistics as calculated with the Z-scores method (**Fig. 2A**), and that rarer cell-types had more extreme Z-score statistics (**Fig. S2** and **S3A-B**). Thus, the links between ATACseq peaks and genes have less significant statistics when identified in the major cell-types of this PBMC dataset.

**Figure 2.**
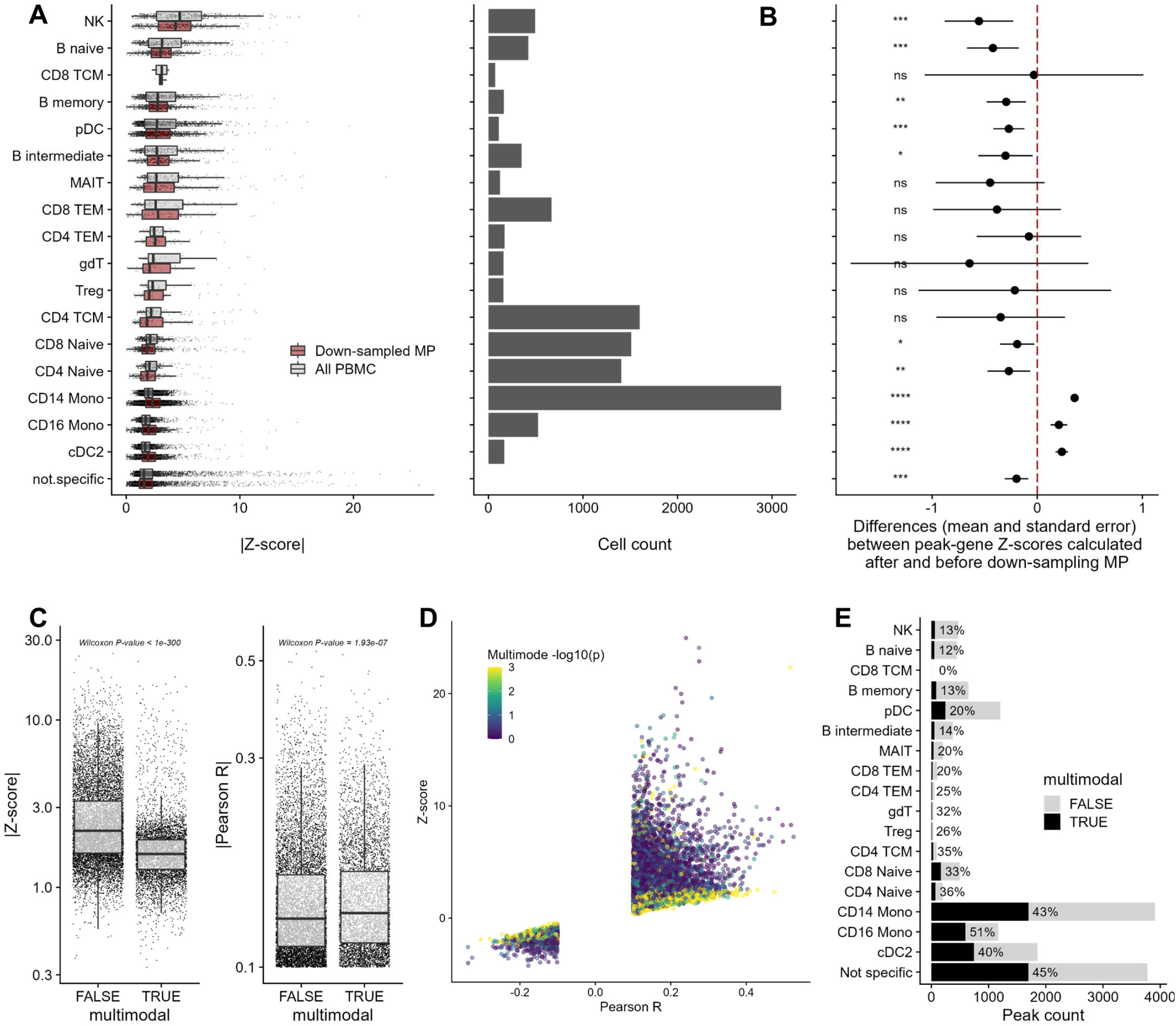
The cell-type composition of the PBMC single-nucleus multiomic dataset impacts the identification of gene-peak links using the Z-scores method. (**A**) Our analyses included 15,113 gene-peak links with |Pearson R| >0.1 (Signac’s default parameter) identified in 30 cell-types. The left column shows the number of cells in each cell-type. In the right column, we show the boxplots of the statistics calculated using the Z-scores method. For this analysis, we assigned gene-peak links to specific cell-type using sensitivity and specificity metrics (**Methods**). Links that could not be unambiguously assigned are grouped in the “not.specific” category. We repeated the analyses by down-sampling the number of mononuclear phagocytes (MP) to n=500. (**B**) Effect of down-sampling mononuclear phagocytes (MP) from 3,788 to 500 cells on peak-gene link statistics calculated with the Z-scores method. Positive values indicate higher Z-scores (i.e. more significant) after down-sampling. ns; not significant, *; P-value<0.05, **; P-value<0.01, ***; P-value<0.001, ****; P-value<0.0001. (**C**) The Z-score statistics and Pearson R coefficients for links between ATACseq peaks and target genes that generated uni- or multimodal null distributions (with the Z-scores method). (**D**) Scatterplot of the Pearson R coefficients (*x*-axis) and statistics calculated with the Z-scores method (*y*-axis) for all links between ATACseq peaks and target genes. Each point is color-coded based on the P-value of the multimode test. Peak-gene links that generated multimode null distributions (in yellow) tend to have Z-score statistics ~ 0 despite many having high Pearson R coefficients. (**E**) Proportion of multimodal null distributions by cell-type generated by the Z-scores method for the tested links between ATACseq peaks and target genes. Mono; Monocytes, cDC; classical Dendritic cells, NK; Natural killer cells, pDC; progenitor Dendritic cells, TEM; T effector memory cells, TCM; T central memory cells, gdT; Gamma delta (γδ) T cells, MAIT; mucosal-associated invariant T cells, Treg; regulatory T cells.

To evaluate if the number of MP influenced the calculated link statistics with the Z-scores method, we down-sampled cells from the MP clusters from 3,788 to 500 cells, and repeated the analyses. The down-sampling increased the Z-scores of cells from the MP clusters (*t*-test P-value_CD14_=1.7×10^−81^, P-value_CD16_=4.5×10^−7^, P-value_cDC2_=3.2×10^−14^), and reduced the peak-gene link Z-scores for all other cell-types except for the ones which had few cell-type-specific ATACseq peaks (**Fig. 2B**). These results suggest that the cell-type composition of the dataset has a strong influence on the statistics calculated using the Z-scores method.

#### More abundant cell-types have more power to identify correlated ATACseq peaks in *trans*

As highlighted for the *NOD2* locus, we found that the Z-scores method often generates bimodal null distributions (**Fig. 1C-D** and **Fig. S4**), and that these bimodal distributions are more frequent in more abundant cell-types (**Fig. S3C**). We hypothesized that the co-accessibility of cell-type-specific *trans*-open chromatin regions – for instance due to the activity of a common transcription factors giving rise to a co-regulatory network – could cause the emergence of a second mode in the null distributions. In support of this hypothesis, removing cell-type-specific *trans*-peaks from the null distributions generally eliminated the bimodality and increased the Z-score statistics (**Fig. 1C-D** and **Fig. S4**). To better understand the effect of the bimodality on the Z-scores method, we tested each null distributions for multiple modes (P-value<0.05 for >1 modes [**Methods**]^10^) and compared the Z-scores and Pearson R coefficients of peak-gene links obtained for multimodal and unimodal null distributions. Whereas statistics from the Z-scores method were significantly lower for peak-gene links with multimodal null distributions (Wilcoxon P-value <1×10^−300^), we found that the simple Pearson R statistics were higher (Wilcoxon P-value=1.93×10^−7^)(**Fig. 2C**). Consistently, we found that peak-gene links with multimodal null distributions were more likely to have non-significant Z-score statistics (near 0), even when the corresponding Pearson R coefficients were relatively high (**Fig. 2D**). Additionally, links between ATACseq peaks and target genes in MP were more likely to have multimodal null distributions when compared to other rarer cell-types (**Fig. 2E**). Together, these results suggest that the popular Z-scores method used to infer a regulatory effect between ATACseq peaks and target genes in multiomic data is biased, counter-intuitively, towards lower abundant cell-types. Our analyses show that this bias arises from the production of bimodal null distributions when matching the tested peak-gene links with links found in *trans* for abundant cell-types, presumably because of increased power to detect co-regulated *trans* ATACseq peaks.

### Read coverage, but not GC content, impacts peak-gene link statistics

Beside the Pearson R and Z-score methods, we considered two additional approaches. First, because of the inherent sparsity of single-nucleus ATACseq data, we tested a zero-inflated negative binomial (ZINB) model, allowing to independently account for the zero component of a peak-gene link. Second, we also tested a new method – scREG – that is reported to outperform the simple Pearson R model on CD14 monocytes peak-gene link predictions based on eQTL data^11,12^. Of note, scREG output link scores for peak-gene within each cell-type. We compared the peak-gene links identified by each of these four models to the Epimap predictions of cCREs and target genes for CD14 monocytes, B cells and NK cells (**Methods**).

In the Z-scores method, the rationale for generating null distributions is to account for possible confounders such as the number of mapped reads (i.e. coverage) and GC bias. We decided to explore the impact of these two factors on the Z-scores, Pearson R, scREG_CD14_, and ZINB statistics. In the Signac workflow, initial filtering is done separately at the gene expression and ATACseq peak level, removing genes or peaks with <10 cells with non-zero counts. Thus, it is possible to have peak-gene links defined by a single cell that has counts for both RNAseq and ATACseq modalities. We found that the Z-scores method was particularly sensitive to the number of cells with non-zero counts, with extreme Z-scores associated with links identified in a small number of cells (**Fig. S5**). Although the same effect was less striking for the Pearson R, scREG_CD14_, and ZINB methods (with high statistics being positively correlated with high number of cells with non-zero counts), we still observed some extreme statistics for links identified in a small number of cells (**Fig. S5**).

We further characterized the impact of GC content on the Z-scores method, as this is the only method that consider this variable. *Trans*-peak matching for an ATACseq peak is done through the attribution of weights dependent on the input variables (here GC and counts). We compared peak-gene link Z-scores from two analyses with identical parameters (i.e. null distributions generated independently twice for the same peak-gene links), and also for analyses with and without GC as a matching criteria. We found that the correlation value of Z-scores for two analyses with identical matching parameters is 0.97 (**Fig. S6A**), while that of a model matching for counts only vs one that matches for both GC content and counts is 0.95 (**Fig. S6B**), suggesting that the GC content does not strongly influence the identification of peak-gene links (even for the stronger links, see the right-hand tail of the distributions in **Fig. S6**).

### The raw Pearson R coefficients and/or physical distance provide better statistics to capture predicted or functionally validated links between ATACseq peaks and target genes

We next turned to independent datasets that have predicted or experimentally ascertained links between cCREs and target genes to address the limitations of the Z-score method and propose new strategies. ZINB is a computationally expensive method compared to other methods tested (**Fig. S7**). Further scREG currently limits its output to 100,000 links. For these reasons, we started by comparing with Epimap predictions the accuracy of the Pearson R and the Z-scores methods when considering a very large number of peak-gene links (|Pearson R| >0.01, n=590,842 links). Our Receiver Operating Characteristics (ROC) curve analysis showed that the Pearson R method outperformed the Z-scores method in these three cell-types (**Fig. 3A**, **S8A** and **S9A**). For instance, the area under the curve (AUC) were 0.71 and 0.51 in CD14 monocytes when applying the Pearson R and Z-scores methods, respectively (**Fig. 3A**). Using a smaller set of peak-gene links with a more stringent threshold for inclusion (|Pearson R| > 0.1, n=15,113 links), we then applied the four methods and compared results with predictions from Epimap. scREG outperformed other models in all cell-types, and the Z-scores method performed worse (**Fig. 3B**, **S8B** and **S9B**).

**Figure 3.**
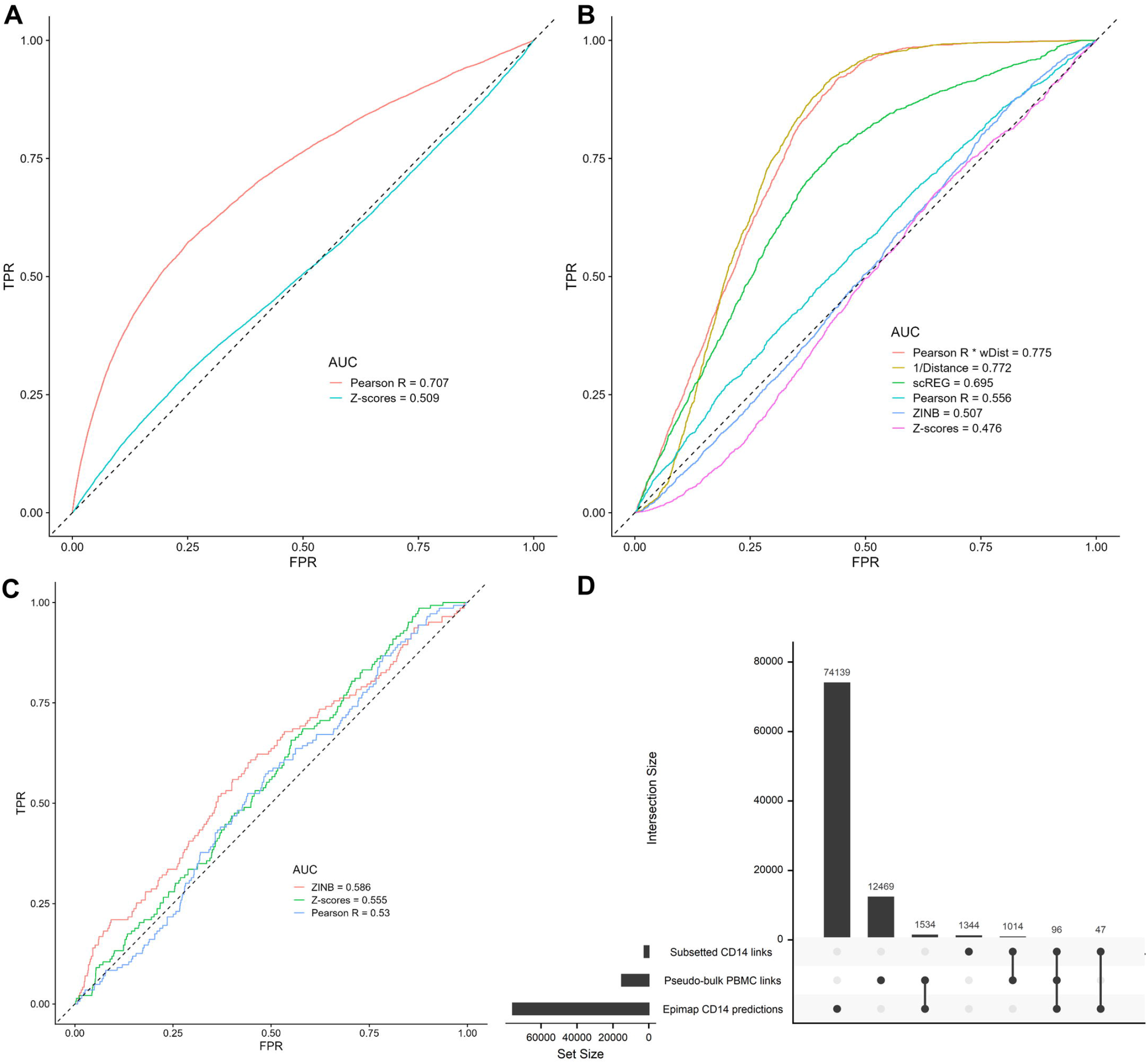
The Pearson R method more accurately validates Epimap-predicted links between cCRE and target genes in CD14 cells. (**A**) We used the Pearson R and Z-scores methods to detect links between ATACseq peaks and target genes (590,842 links with |Pearson R| >0.01) in the complete (i.e. pseudo-bulk) PBMC multiomic dataset. Then, we performed Receiving Operating Curves (ROC) analyses to compare the identified peak-gene links from the multiomic data with regulatory links in CD14 cells predicted by the Epimap Project. (**B**) As in **A**, but using a smaller set of links defined using a more stringent statistical threshold (15,113 links with |Pearson R| >0.1). All cell-types are used to identify links (pseudo-bulk), except for scREG which by design output link scores by cell-type (in this case, CD14 cells). (**C**) As in **B**, but limiting these ROC analyses to links between ATACseq peaks and target genes with |Pearson R| >0.1 that were found in the CD14 cells subset of the PBMC multiomic dataset. (**D**) Upset plot that shows the intersections of links identified between ATACseq peaks and target genes using either the full PBMC multiomic dataset (i.e. pseudo-bulk) or only the CD14 cells subset with cCRE-gene regulatory links in CD14 cells as predicted by the Epimap Project. ZINB; zero-inflated negative binomial, wDist; weighted distance (e^(-distance/200kb)^), TPR, true positive rate; FPR, false positive rate.

The physical distance between regulatory sequences and gene transcription start sites has been found to be a strong predictor of cCRE’s effects on nearby genes^3,13^. Because the scREG model weights the peak-gene link scores with physical distance (e^(-distance/200kb)^), we reasoned that weighting the Pearson R coefficients by the distance between the ATACseq peaks and the target genes could improve its accuracy. Physical distance-weighted Pearson R coefficients resulted in AUC that were similar to those obtained when using distance alone, and remarkedly better than with scREG on all three Epimap cell-type predictions (**Fig. 3B**, **S8B** and **S9B**).

As described above, scREG calculates peak-gene link scores per cell-type^12^. We repeated the analyses of the Epimap predictions with the Z-scores, Pearson R and ZINB methods but focusing on single cell-type. For instance, for the Epimap CD14 predictions, we only analyzed peak-gene links identified in the CD14 subset of the PBMC multiomic dataset. This approach had a minimal impact on the AUC statistics for all three cell-types analyzed (compare panels **B** and **C** in **Fig. 3** (CD14 cells), **Fig. S8** (B cells) and **Fig. S9** (NK cells)), but severely reduced the number of detected peak-gene links (**Fig. 3D**, **Fig. S8D** and **Fig. S9D**). One major drawback of this single cell-type approach is that by reducing the number of cells in the analyses, we significantly reduced power to detect peak-gene links. For instance, our analysis of the whole PBMC dataset (i.e. pseudo-bulk) yielded 15,113 peak-gene links with |Pearson R| > 0.1, including 1611 links (10.7%) that overlap with Epimap predictions for CD14 cells. In contrast, when we restricted our analysis to CD14 cells from the PBMC multiomic dataset, we found 2,499 links, including 143 (5.7%) also predicted by Epimap in CD14 cells (**Fig. 3D**). From these observations, we conclude that using all available cells from multiomic experiments to detect links between ATACseq peaks and target genes is a more powerful strategy than limiting the analyses to single cell-type.

Finally, we compared the peak-gene links identified in the PBMC multiomic datasets with 644 cCRE-gene pairs that were functionally validated using CRISPR perturbations in different cell models^14^. We found that distance alone (or in combination with the Pearson R coefficient) was the best predictor of links between ATACseq peaks and genes that were consistently validated by CRISPR perturbations (**Fig. 4**). We also noted that the simple Pearson R statistic, even without being weighted by physical distance (AUC=0.73), outperformed all other metrics, including the distance-weighted scREG scores (AUC=0.65-0.67)(**Fig. 4**).

**Figure 4.**
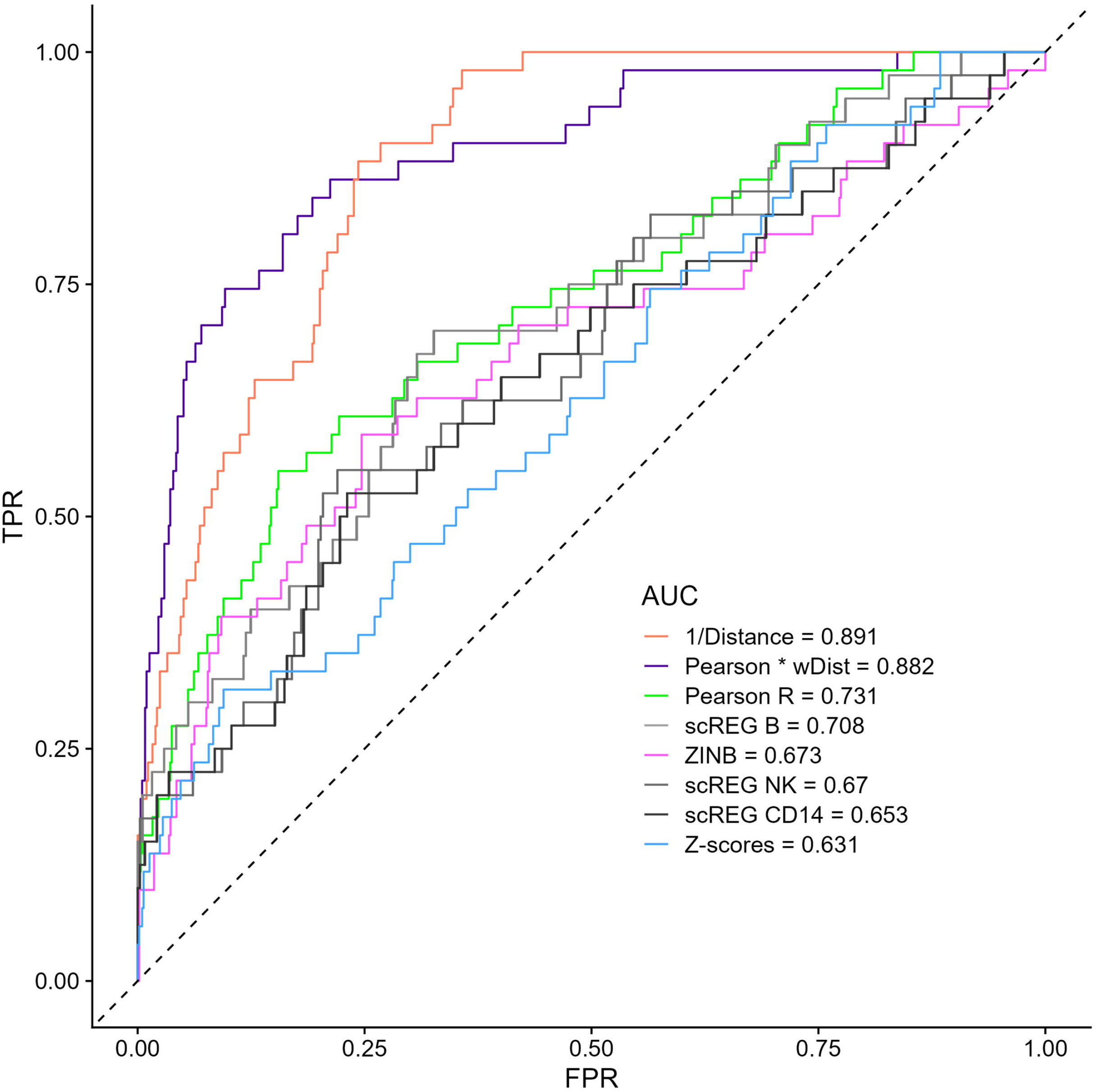
Physical distance and the Pearson R coefficient best capture cCRE-gene pairs identified by CRISPR perturbations. We identified 644 CRISPR-validated cCRE-gene pairs that had corresponding links (defined using |Pearson R| > 0.01) in the PBMC multiomic dataset. Distance-alone or distance-weighted Pearson R coefficients are the best predictors, with the Z-score method (implemented in Signac) performing worst. ZINB; zero-inflated negative binomial, wDist; weighted distance (e^(-distance/200kb)^), TPR, true positive rate; FPR, false positive rate.

## Discussion

Motivated by the absence of several strong candidate links between regulatory sequences and target genes in our analysis of multiomic PBMC data using a common bioinformatic pipeline, we investigated several factors that could impact these results. We found that cell-type composition in single-nucleus data can have a dramatic effect on the ability to detect peak-gene links. Indeed, we showed that null distributions matched on ATACseq peak coverage and GC content are often bimodal, especially when the peaks are specific to (or enriched in) the major cell-types. Our analyses suggest that this second mode arise because of following reasons: First, the number of ATACseq peaks detected in a given cell-type increases with the number of cells, thus increasing the chances to draw *trans*-ATACseq peaks that are opened in that cell-type when building the null distributions. Second, as cells within a given cell-type share transcription factors, their open chromatin regions tend to also be more correlated. Together, this creates two modes: one coming from the *trans*-ATACseq peaks of the dominant cell-type and a second from other cell-types (generally less correlated). We illustrate this conclusion by showing that if a peak-gene link is cell-type-specific, removing ATACseq peaks detected in this cell-type from the null distribution generally removes the mode most associated with the tested link (**Fig. 1C-D**) and drastically increases its Z-score (**Fig. 2B**). This approach also has the consequence to inflate the Z-scores of links mostly found in less abundant cell-types, because the null distributions will contain fewer ATACseq peaks from the same, rare cell-types (**Fig. 2B**). One apparent solution to this problem is to detect peak-gene links within specific cell-types, although we showed that this method is sub-optimal because of the loss in power to detect peaks and genes when fewer cells are analyzed.

The rationale behind the null distributions implemented in the Z-scores model comes from reports that the Tn5 transposase used in the ATACseq protocol has a GC bias^15^. However, a recent comprehensive study specifically addressing this issue did not detect such bias^16^. Furthermore, our own analyses found minimal (if any) effect of GC content the detection of links between ATACseq peaks and target genes, while the number of cells with non-zero counts in both the ATACseq peak and the gene is a major determinant. We recommend not analyzing links if at least 5 cells do not have both non-zero counts in the ATACseq and RNAseq modalities.

There are no perfect datasets to validate links between ATACseq peaks and the promoter of target genes that are inferred from single-nucleus multiomic experiments. In our study, we used predictions from Epimap and published perturbations using CRISPR tools. Surprisingly, we found that simply considering physical distance and/or the Pearson correlation coefficients provide optimal concordance with these datasets. These methods also have the advantage to be computationally scalable, something that can become problematic for the ZINB method (and to some extent the scREG and Z-scores methods as well). It is obvious that larger multiomic experiments as well as true “gold-standard” datasets of *bona fide* peak-gene links will enable the development of more sophisticated statistical methods. In the meantime, we recommend to carefully consider “ATACseq peaks-target genes” links inferred from single-nucleus multiomic analyses, and to validate them using orthogonal approaches such as 3D chromatin conformation analyses, expression quantitative trait loci (eQTL) results, and *in silico* predictions.

## Methods

### Multiomic PBMC data

We analyzed the PBMC multiomic dataset from 10X Genomics (https://www.10xgenomics.com/resources/datasets/pbmc-from-a-healthy-donor-granulocytes-removed-through-cell-sorting-10-k-1-standard-1-0-0). The data was processed according to the Signac tutorial (https://satijalab.org/signac/articles/pbmc_multiomic.html), which uses the same dataset. For the 11,331 cells identified, the workflow annotated 29 cell-types, of which 17 have more than 50 cells (cell-types represented in the dendrogram in **Fig. S1A**). We restricted our analyses to those 17 cell-types for power purposes. For all *cis*-links, we only analyzed ATACseq peaks located within 500kb of a gene transcription start site.

### Cell-type marker ATACseq peaks

The accessibility specificity of ATACseq peaks was tested using the Presto package (*wilcoxauc.Seurat()* function). A marker ATACseq peak was attributed to a cell-type using the highest area under the curve (AUC). ATACseq peaks with AUCs < 0.55 and an FDR > 10^−5^ in all cell-types with more than 50 cells were attributed to the non-specific peak group.

### Down-sampling mononuclear phagocytes

Cell-types clustering together, both in RNAseq and ATACseq UMAPs, were categorised as part of mononuclear phagocytes (MP)(CD14 Mono, CD16 Mono, cDC2) cluster together in both the RNAseq and ATACseq UMAPs. 500 MP out of 3,782 cells were randomly drawn and reprocessed with the rest of the PBMC cells. We compared Z-scores of peak-gene links with overlapping peaks and identical genes between the full dataset (n=11,331 cells) and the down-sampled MP dataset (n =8,049 cells).

### Removing co-regulated peaks from null distributions

We used the output of Signac’s *CallPeaks()* function to retrieve from which cell-type a peak was called by MACS2^17^ (implemented in Signac). The cell-types were categorised in 4 broader classes representative of the UMAP and dendrogram:

1. **Lymphoid**; CD8 Naive, CD4 Naive, CD4 TCM, CD8 TEM, CD8 TCM, CD4 TEM, MAIT, Treg
2. **NK cells**; gdT, NK, CD8 TEM, MAIT
3. **Monocytes**; CD14 Mono, CD16 Mono, cDC2, pDC
4. **B cells**; B intermediate, B memory, B naive

For ATACseq peaks that were called in all 4 classes, no filtering was done. For ATACseq peaks with some specificity, we removed all *trans*-peaks also called in the same broad cell-type class. Therefore, a tested ATACseq peaks called in B cells and Monocytes by MACS2, but not called in lymphoid and NK cells would have a null distribution composed of *trans*-peaks called in lymphoid and NK cells.

### Multimodal test

To establish that a null distribution was multimodal, we used the mixtools package expectation–maximization function *normalmixEM()* with k=2 and epsilon = 1e-03 as described in Ameijeiras-Alonso *et al*., 2019^18^. To better assess the bimodality of null distributions, we increased the number of *trans*-ATACseq peaks used to 1000. Null distributions with nominal p-values < 0.05 were categorised as multimodal.

### Pearson R and Z-score models

#### Pseudo-bulk

We used the R package Signac function’s *LinkPeaks()* with a 500kb window for a gene’s transcription starting site and a null distribution of 200 *trans*-ATACseq peaks to obtain Z-scores and Pearson R as described in the package tutorial (https://satijalab.org/signac/articles/pbmc_multiomic.html) on all PBMC passing the quality-control steps. We used the log(ATACseq peak sum of counts +1) instead of the counts to match peaks. Peak-gene links were filtered for |Pearson R| > 0.01 to remove cells with zero count in both the ATACseq peak and the gene tested or > 0.1 to limit the number of tests as mentioned in the text and the figure legends.

#### Cell-type subsetted

The same strategy mentioned in *<u>Pseudo-bulk</u>*, was applied independently after subsetting the CD14 mono cells, B cells (including B intermediate, B memory, B naive) and NK cells (NK, NK Proliferating, NK_CD56bright).

### ZINB model

We tested the ZINB model below using the R package pscl function *zeroinfl()*^19^.

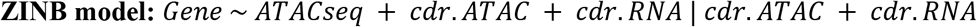

The zero-inflated component was modeled with cellular detection rates (cdr) for both ATACseq peaks and genes (proportion of features with 0 counts). For genes with no 0 counts across all cells, we used the negative binomial generalized linear (GLMNB) model implemented with the R package MASS function *glm.nb()* given that no zero component could be modeled.

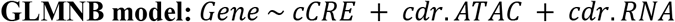

The |Z-value| were used as predictive value for each tested peak-gene links.

### scREG implementation

We initially tested an exact implementation of the package tutorial (https://github.com/Durenlab/RegNMF). The current software returns a prioritized list of links (10,000 per identified clusters) with very low overlap with our validation data, which made the comparison with Epimap and CRISPRi validation uninformative. We found 2 likely explanations. First, the output from *SplitGroup()* are the 10,000 peak-gene pairs with lowest scores values for that cell type, as opposed to the 10,000 highest scores. Second, the ATACseq data is log10 transformed while the RNAseq data is log2 transformed. This creates stronger weights for the gene expression component of the links matrix and skews results towards highly expressed genes (i.e. ATACseq peaks linked to *MALAT1* were the top 20 links for all clusters). To have comparable results to the other tested models, we used as inputs the genes and peaks from the 15,113 peak-gene links with |Pearson R| > 0.1 as well as the 644 peak-gene links kept from CRISPR validations.

### Peak-gene link models comparison with Epimap

We retrieved the peak-gene link predictions for the PBMC matching cell-types (CD14 MONOCYTE, B CELL, NK CELL) from https://personal.broadinstitute.org/cboix/epimap/links/links_corr_only/. We chose these cell-types because their cluster showed greater homogeneity and boundaries (in contrast to the lymphoid cells, see **Fig. S1**). For each cell-type, we kept peak-gene links found in all replicates to insure reproducibility. We recovered the hg38 positions using the AnnotationHub package (Annotationhub chain: hg19ToHg38.over.chain.gz). For these analyses, cell-type-specific peak-gene links from the PBMC multiomic dataset were considered true positive when the ATACseq peak overlapped at least partly the enhancer position and the linked gene was the same. ROC curves were calculated with the ROCR package by increasing the thresholds of the model’s statistic.

### Peak-gene links validation with CRISPR perturbation results

Given the modest number of CRISPR-based validated links, we expanded the number of tested links to include all links with non-zero read counts for both the gene and the ATACseq peak tested (|Pearson R| > 0.01 (n=590,842)). ROC curves were calculated using the ROCR package in a set of 644 CRISPR validations from *J. Nasser et al. 2021* (Table S5) overlapping PBMC links. The original CRISPR validation data (n=5755) was filtered to include CRISPR targets overlapping an ATACseq peak with a corresponding gene expression readout. We also excluded duplicates (same link tested in multiple cell lines), links that showed divergent results across cell lines and those that were excluded by the author of the study for various reasons, denoted by the *IncludeInModel* column (power insufficient, overlapping promotor and others).

## Data availability

All results presented here were generated with our code available on GitHub: https://github.com/lebf3/Links_models_multomic.

## Acknowledgements

This work was funded by the Canadian Institutes of Health Research (MOP #136979), the Canada Research Chair Program, the Foundation Joseph C. Edwards and the Montreal Heart Institute Foundation (G. Lettre). F. JA Leblanc was supported by the Fonds de Recherche en Santé du Québec (FRQS) and Université de Montréal.

## Notes

https://satijalab.org/signac/articles/pbmc_multiomic.html

https://personal.broadinstitute.org/cboix/epimap/metadata/Imputation_Metadata.xlsx

https://static-content.springer.com/esm/art%3A10.1038%2Fs41586-021-03446-x/MediaObjects/41586_2021_3446_MOESM7_ESM.txt

### Competing Interest Statement

The authors have declared no competing interest.

