## Supplementary Figures for "Major cell-types in multiomic single-nucleus datasets impact statistical modeling of links between regulatory sequences and target genes"

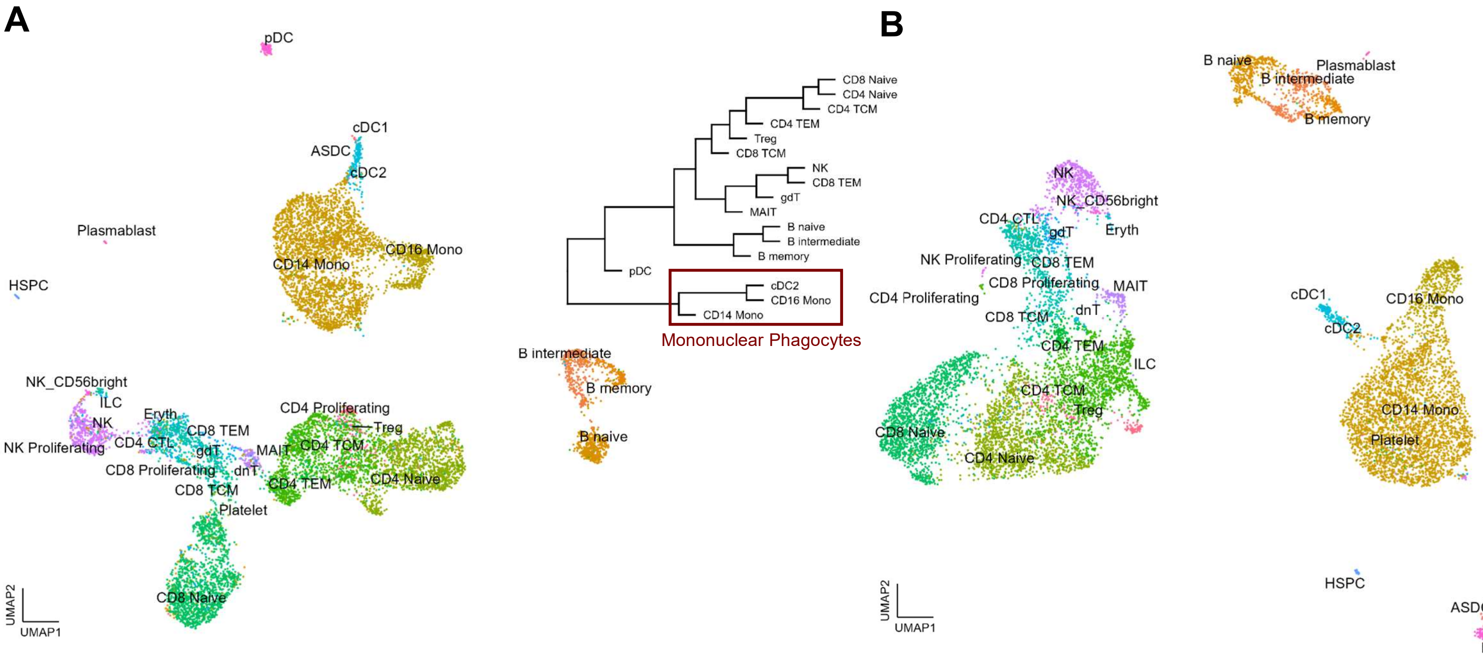

A
pDC
cDC1
ASDC
cDC2
Plasmablast
CD16 Mono
CD14 Mono
HSPC
NK_CD56bright
ILC
Eryth
NK
CD8 TEM
CD4 Proliferating
Treg
NK Proliferating
CD4 CTL
gdT
MAIT
CD4 TCM
CD8 Proliferating
dnT
CD8 TCM
CD4 TEM
CD4 Naive
Platelet
CD8 Naive
UMAP2
UMAP1
CD8 Naive
CD4 Naive
CD4 TCM
CD4 TEM
Treg
CD8 TCM
NK
CD8 TEM
gdT
MAIT
B naive
B intermediate
B memory
pDC
cDC2
CD16 Mono
CD14 Mono
Mononuclear Phagocytes
B intermediate
B memory
B naive
B
B naive
Plasmablast
B intermediate
B memory
NK
NK_CD56bright
CD4 CTL
gdT
Eryth
NK Proliferating
CD8 TEM
CD8 Proliferating
MAIT
CD4 Proliferating
CD8 TCM
dnT
CD4 TEM
ILC
CD4 TCM
Treg
CD8 Naive
CD4 Naive
cDC1
CD16 Mono
cDC2
CD14 Mono
Platelet
HSPC
ASDC
pDC
UMAP2
UMAP1

**Supplementary Figure 1**. Clustering of 11,331 PBMC using (**A**) ATACseq and (**B**) RNAseq data. The data is embedded using uniform manifold approximation and projection (UMAP). The Euclidean distance of mean expression by cell-type represented as dendrogram shows mononuclear phagocytes as a distinct cell archetype (CD14, CD16 and cDC2).

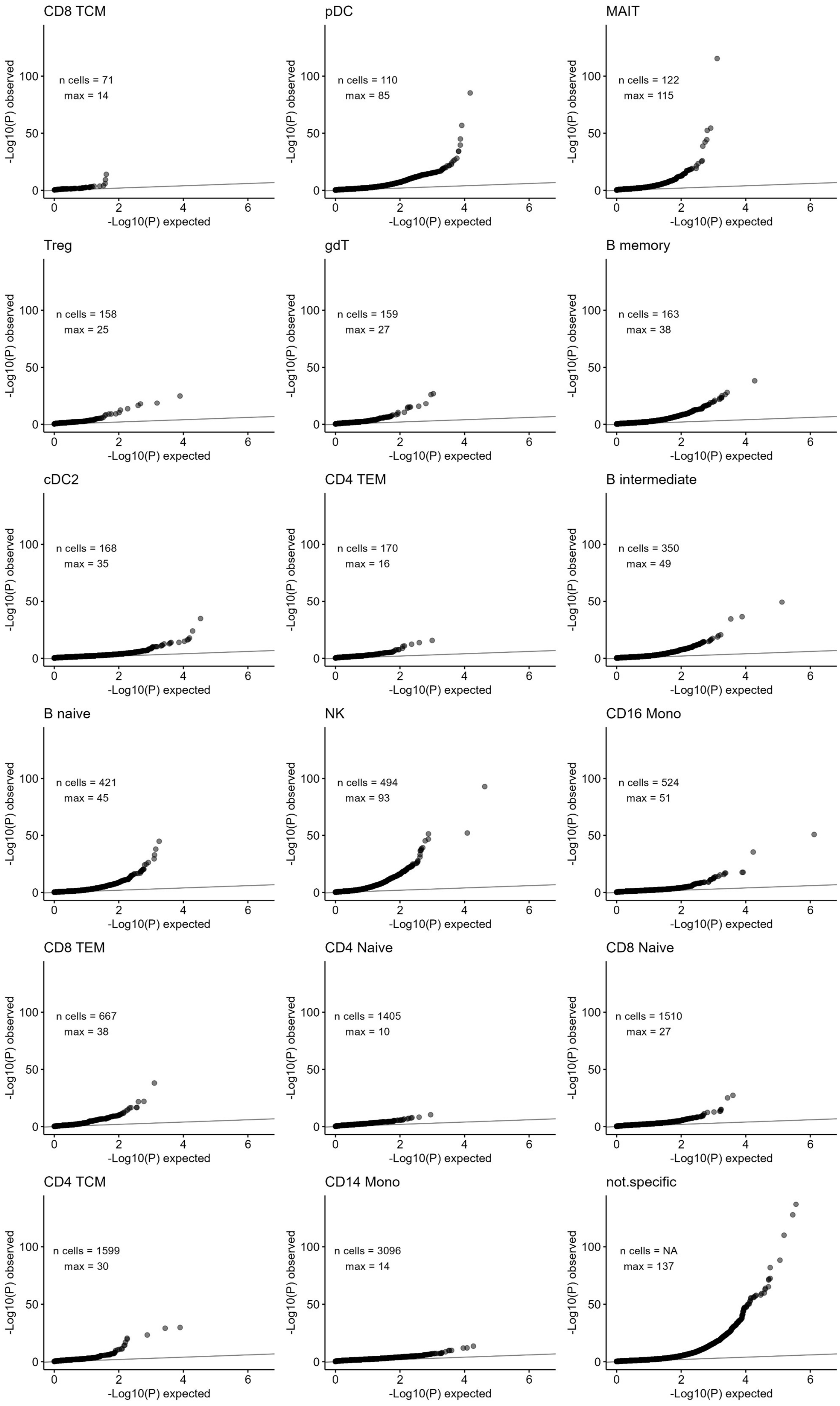
CD8 TCM
n cells = 71
max = 14
pDC
n cells = 110
max = 85
MAIT
n cells = 122
max = 115
Treg
n cells = 158
max = 25
gdT
n cells = 159
max = 27
B memory
n cells = 163
max = 38
cDC2
n cells = 168
max = 35
CD4 TEM
n cells = 170
max = 16
B intermediate
n cells = 350
max = 49
B naive
n cells = 421
max = 45
NK
n cells = 494
max = 93
CD16 Mono
n cells = 524
max = 51
CD8 TEM
n cells = 667
max = 38
CD4 Naive
n cells = 1405
max = 10
CD8 Naive
n cells = 1510
max = 27
CD4 TCM
n cells = 1599
max = 30
CD14 Mono
n cells = 3096
max = 14
not.specific
n cells = NA
max = 137
-Log10(P) observed
-Log10(P) expected

**Supplementary Figure 2**. Distributions of ATACseq peaks-gene link statistics calculated using the Z-scores method as implemented in Signac. The peak-gene links with |Pearson R > 0.01| were attributed to specific cell-type (cell-types with n>50 cells) using the peak's specificity of accessibility (**Methods**). Mono; Monocytes, cDC; classical Dendritic cells, NK; Natural killer cells, pDC; progenitor Dendritic cells, TEM; T effector memory cells, TCM; T central memory cells, gdT; Gamma delta (γδ) T cells, MAIT; mucosal-associated invariant T cells, Treg; regulatory T cells, Max; maximal $-\log_{10}$(P-value) calculated for a given cell-type.

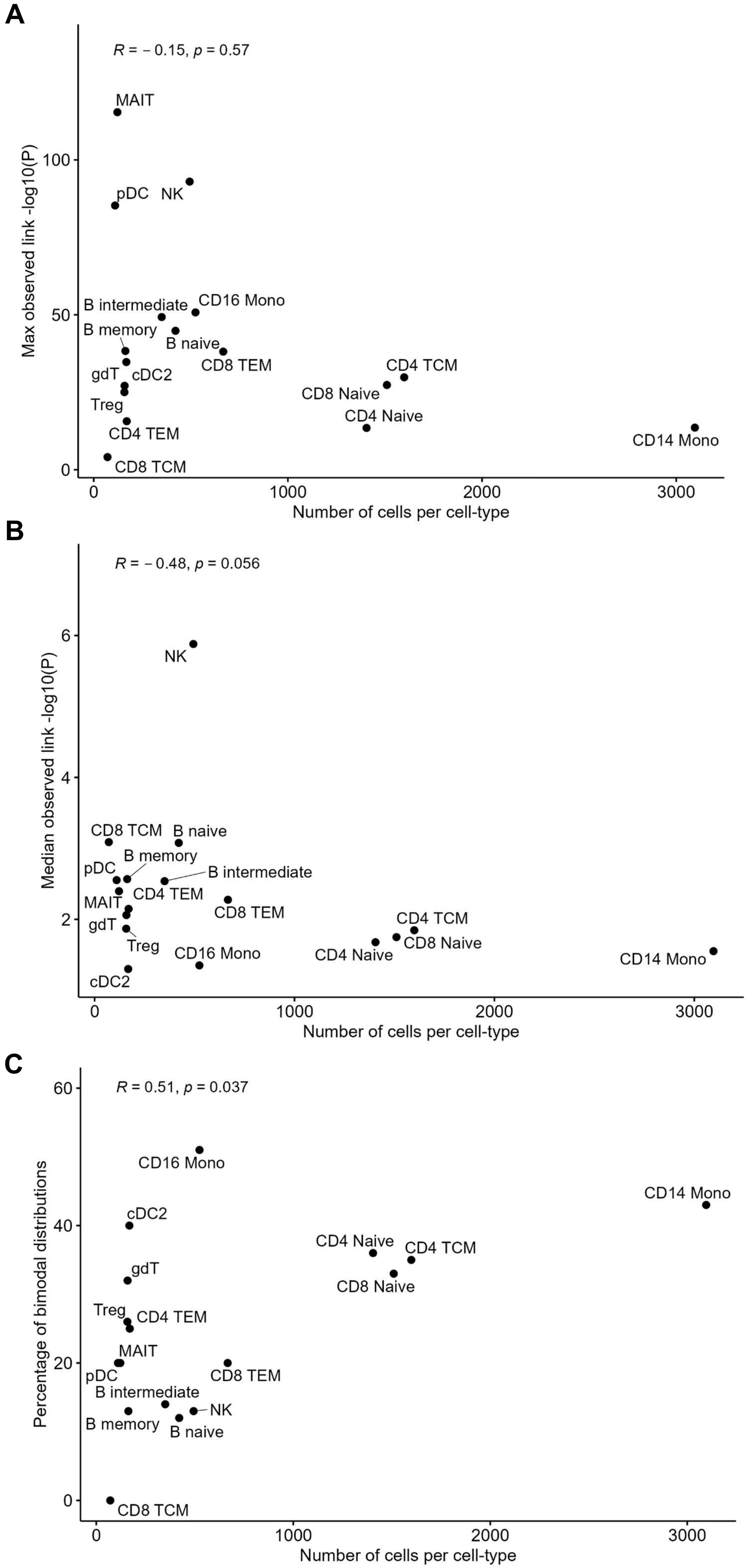
A
R = − 0.15, p = 0.57
Max observed link -log10(P)
Number of cells per cell-type
MAIT
pDC
NK
B intermediate
CD16 Mono
B memory
B naive
CD8 TEM
gdT
cDC2
Treg
CD4 TEM
CD8 TCM
CD4 TCM
CD8 Naive
CD4 Naive
CD14 Mono
0
50
100
0
1000
2000
3000
B
R = − 0.48, p = 0.056
Median observed link -log10(P)
Number of cells per cell-type
NK
CD8 TCM
B naive
B memory
pDC
B intermediate
MAIT
CD4 TEM
CD8 TEM
gdT
Treg
CD16 Mono
cDC2
CD4 TCM
CD8 Naive
CD4 Naive
CD14 Mono
2
4
6
0
1000
2000
3000
C
R = 0.51, p = 0.037
Percentage of bimodal distributions
Number of cells per cell-type
CD16 Mono
CD14 Mono
cDC2
CD4 Naive
CD4 TCM
gdT
CD8 Naive
Treg
CD4 TEM
MAIT
pDC
CD8 TEM
B intermediate
NK
B memory
B naive
CD8 TCM
0
20
40
60
0
1000
2000
3000

**Supplementary Figure 3**. The impact of cell-type counts on properties of the Z-scores method implemented in Signac. For cell-type-specific ATACseq peaks, we identified peak-gene links and compared (**A**) the most extreme P-value from the Z-score method, (**B**) the median P-value from the Z-score method, or (**C**) the percentage of bimodal null distributions with the number of cells in that cell-type. P-values are from the Spearman correlation test. Mono; Monocytes, cDC; classical Dendritic cells, NK; Natural killer cells, pDC; progenitor Dendritic cells, TEM; T effector memory cells, TCM; T central memory cells, gdT; Gamma delta (γδ) T cells, MAIT; mucosal-associated invariant T cells, Treg; regulatory T cells.

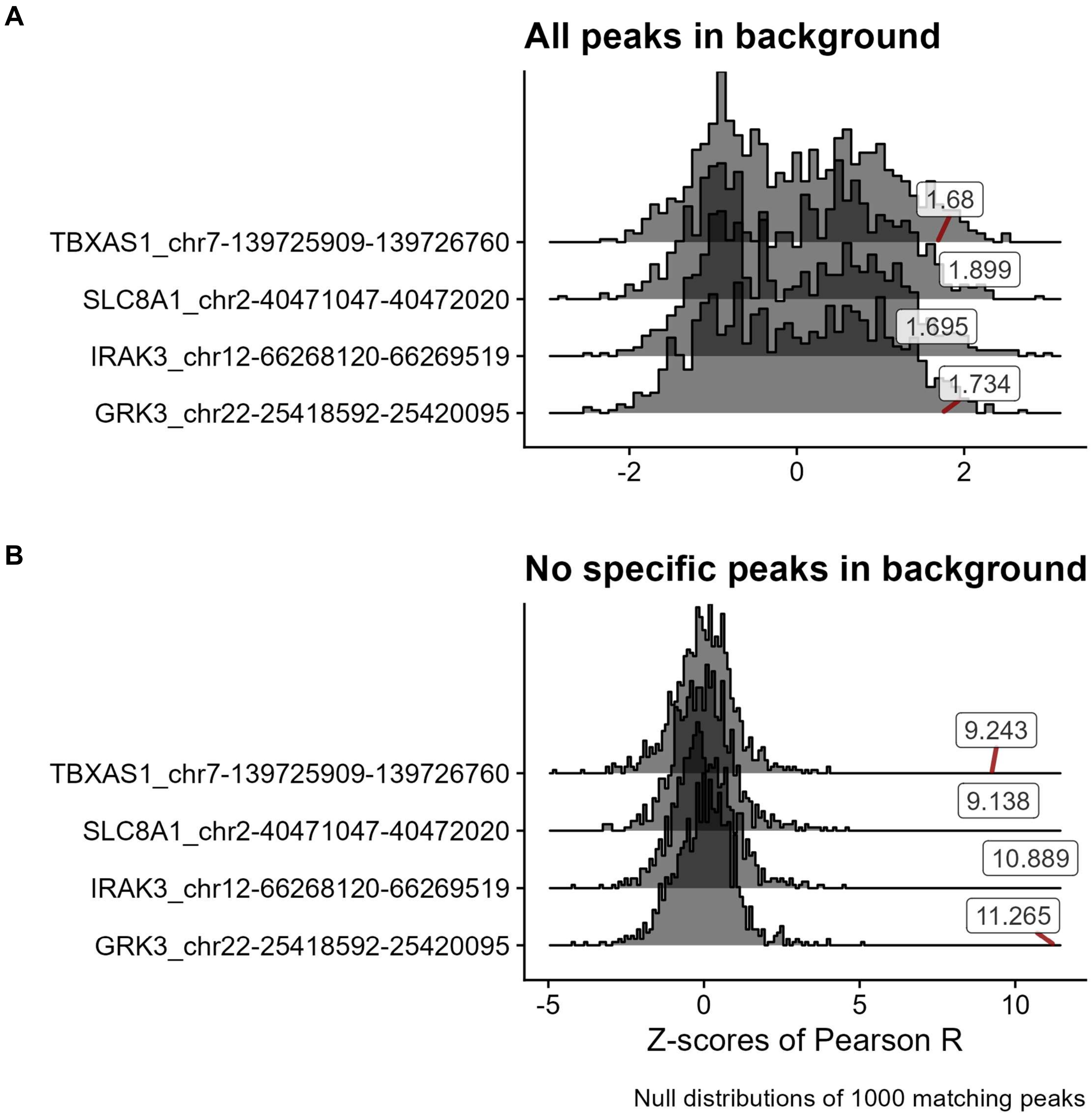
A
All peaks in background
TBXAS1_chr7-139725909-139726760
SLC8A1_chr2-40471047-40472020
IRAK3_chr12-66268120-66269519
GRK3_chr22-25418592-25420095
1.68
1.899
1.695
1.734
-2
0
2
B
No specific peaks in background
TBXAS1_chr7-139725909-139726760
SLC8A1_chr2-40471047-40472020
IRAK3_chr12-66268120-66269519
GRK3_chr22-25418592-25420095
9.243
9.138
10.889
11.265
-5
0
5
10
Z-scores of Pearson R
Null distributions of 1000 matching peaks

**Supplementary Figure 4**. Other examples of bimodal null distributions generated by the Z-scores method. Four GC- and count-matched null distributions with high Pearson R coefficients and low Z-scores. Labeled boxes represent the Z-scores for the tested cCRE and its linked gene (*y*-axis names: gene_peak) against the null distributions (**A**) before and (**B**) after removing from the dataset *trans* ATACseq peaks that are specific to the cell-type in which the cCRE is mostly accessible.

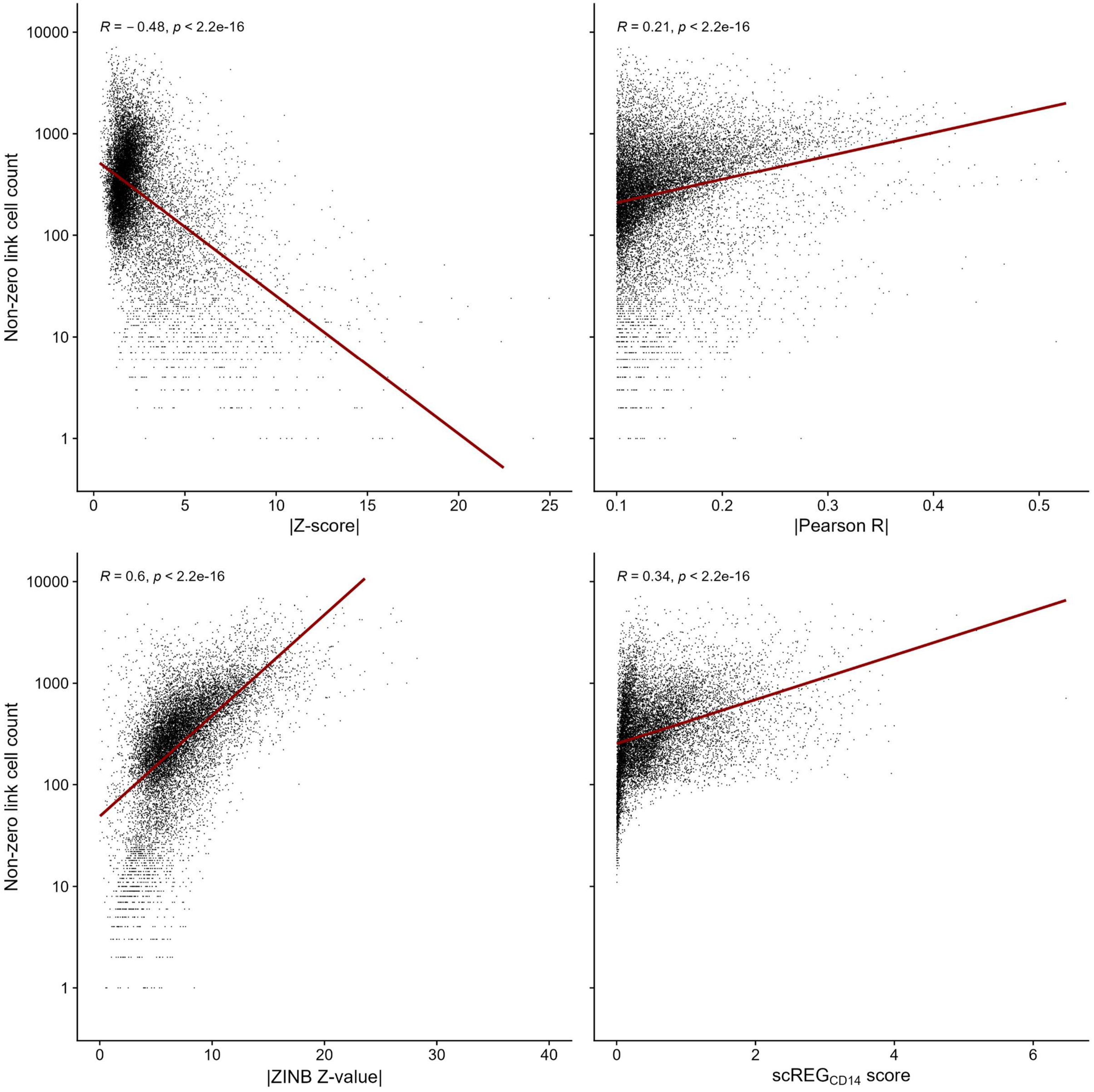

R = − 0.48, p < 2.2e-16
R = 0.21, p < 2.2e-16
R = 0.6, p < 2.2e-16
R = 0.34, p < 2.2e-16
Non-zero link cell count
|Z-score|
|Pearson R|
|ZINB Z-value|
scREG CD14 score

**Supplementary Figure 5**. The Z-scores method tends to output extreme statistics for peak-gene links that are identified in a few cells. In contrast, the statistics for the Pearson R, ZINB and $scREG_{CD14}$ methods are higher when there are more cells with non-zero counts (as expected given higher power to detect links). Z-scores, ZINB Z-values and Pearson R from links with |Preason R| > 0.1 were compared against the number of cells for which both the gene and the peak from that link had a non-zero read count. We found one outlier using ZINB which was removed for visualisation. On each plot, we added the Pearson R coefficient and corresponding P-value.

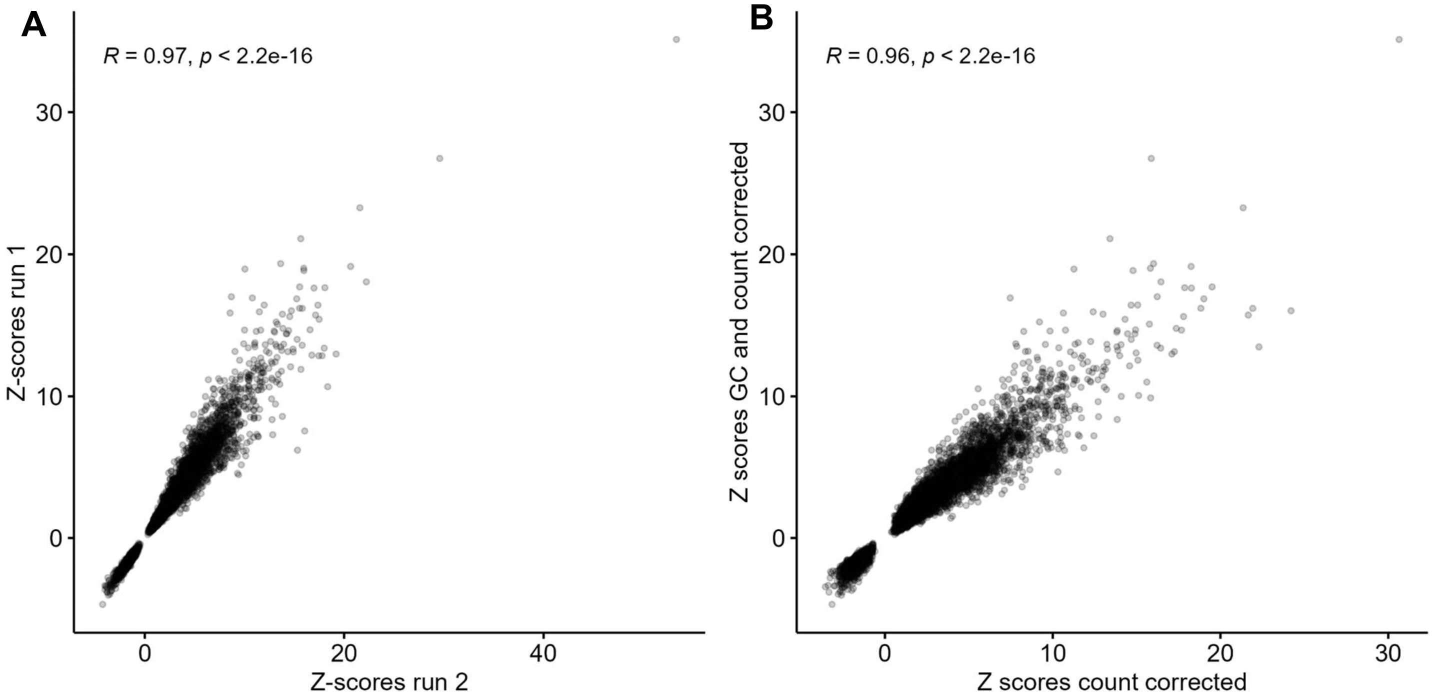
A
R = 0.97, p < 2.2e-16
Z-scores run 1
Z-scores run 2
0
10
20
30
0
20
40
B
R = 0.96, p < 2.2e-16
Z scores GC and count corrected
Z scores count corrected
0
10
20
30
0
10
20
30

**Supplementary Figure 6**. Accounting for GC content has minimal impact on the peak-gene link statistics. (**A**) Distribution of Z-scores for 2 analyses of all peak-gene links with |Pearson R| > 0.1 using the same model. The variability is due to the stochastic sampling of peaks to create null distributions. (**B**) Distribution of statistics comparing the Z-scores model matching peaks for both GC percent and counts (*y*-axis) or counts only (*x*-axis). On each plot, we added the Pearson R coefficient and corresponding P-value.

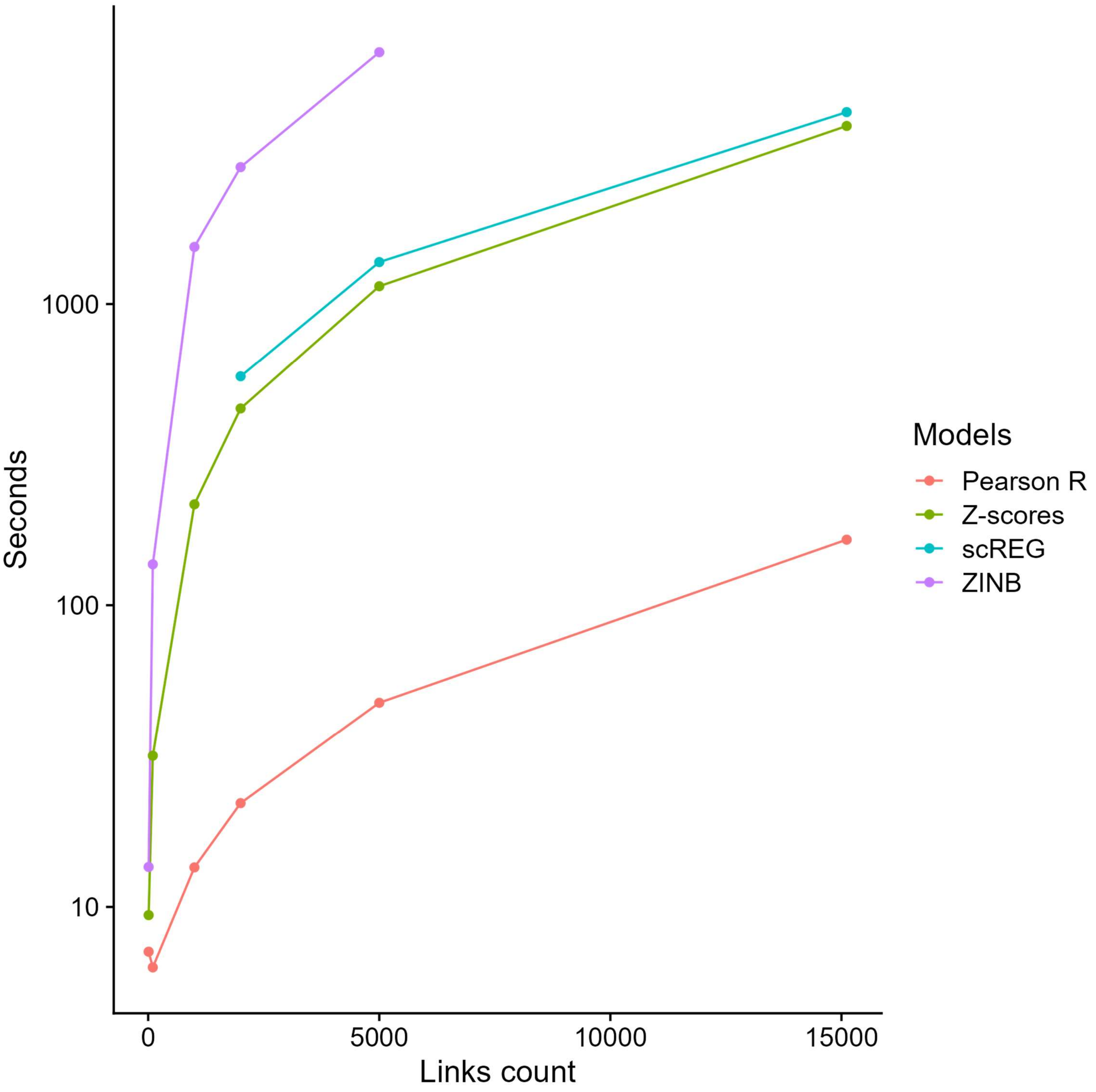
Seconds
1000
100
10
0
5000
10000
15000
Links count
Models
Pearson R
Z-scores
scREG
ZINB

**Supplementary Figure 7**. The Pearson R provides an important scalability advantage. We benchmarked times to run each of the 4 models tested using 1 core with an AMD Ryzen 7 5800X 3,8GHz processor. For each model we tested 10, 100, 1000, 5000 and 15000 links except for ZINB which shows poor scalability. scREG returned errors for inputs with < 2000 links.

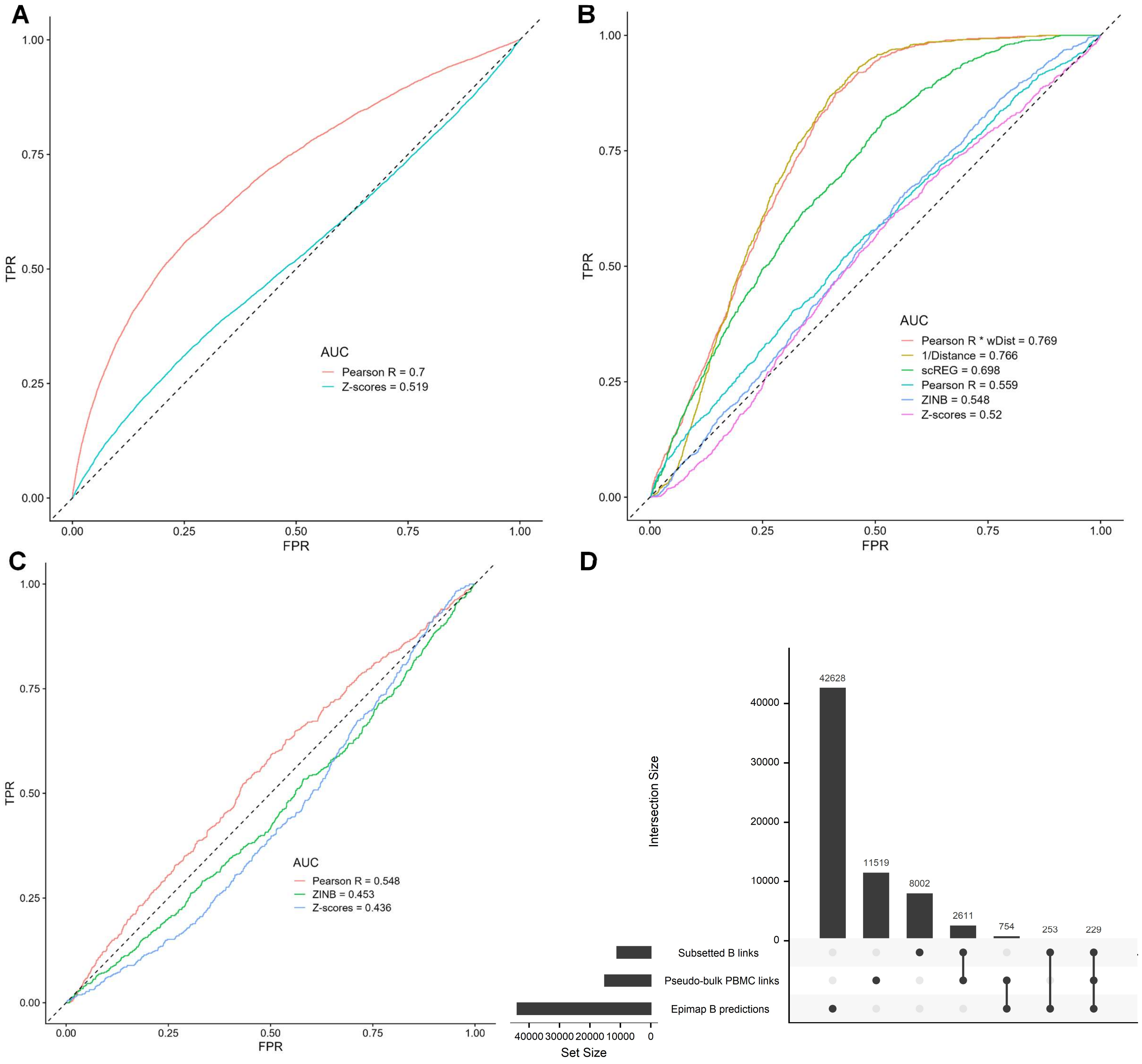
A
TPR
FPR
0.00
0.25
0.50
0.75
1.00
AUC
Pearson R = 0.7
Z-scores = 0.519
B
AUC
Pearson R * wDist = 0.769
1/Distance = 0.766
scREG = 0.698
Pearson R = 0.559
ZINB = 0.548
Z-scores = 0.52
C
AUC
Pearson R = 0.548
ZINB = 0.453
Z-scores = 0.436
D
Intersection Size
42628
11519
8002
2611
754
253
229
Subsetted B links
Pseudo-bulk PBMC links
Epimap B predictions
Set Size
40000
30000
20000
10000
0

**Supplementary Figure 8**. The Pearson R method more accurately validates Epimap-predicted links between cCRE and target genes in B cells. (**A**) We used the Pearson R and Z-scores methods to detect links between ATACseq peaks and target genes (590,842 links with |Pearson R| >0.01) in the complete (i.e. pseudo-bulk) PBMC multiomic dataset. Then, we performed Receiving Operating Curves (ROC) analyses to compare the identified peak-gene links from the multiomic data with regulatory links in B cells predicted by the Epimap Project. (**B**) As in **A**, but using a smaller set of links defined using a more stringent statistical threshold (15,113 links with |Pearson R| >0.1). All cell-types are used to identify links (pseudo-bulk), except for scREG which by design output link scores by cell-type (in this case, B cells). (**C**) As in **B**, but limiting these ROC analyses to links between ATACseq peaks and target genes with |Pearson R| >0.1 that were found in the B cells subset of the PBMC multiomic dataset. (**D**) Upset plot that shows the intersections of links identified between ATACseq peaks and target genes using either the full PBMC multiomic dataset (i.e. pseudo-bulk) or only the B cells subset with cCRE-gene regulatory links in B cells as predicted by the Epimap Project. ZINB; zero-inflated negative binomial, wDist; weighted distance ($e^{(-distance/200kb)}$), TPR, true positive rate; FPR, false positive rate.

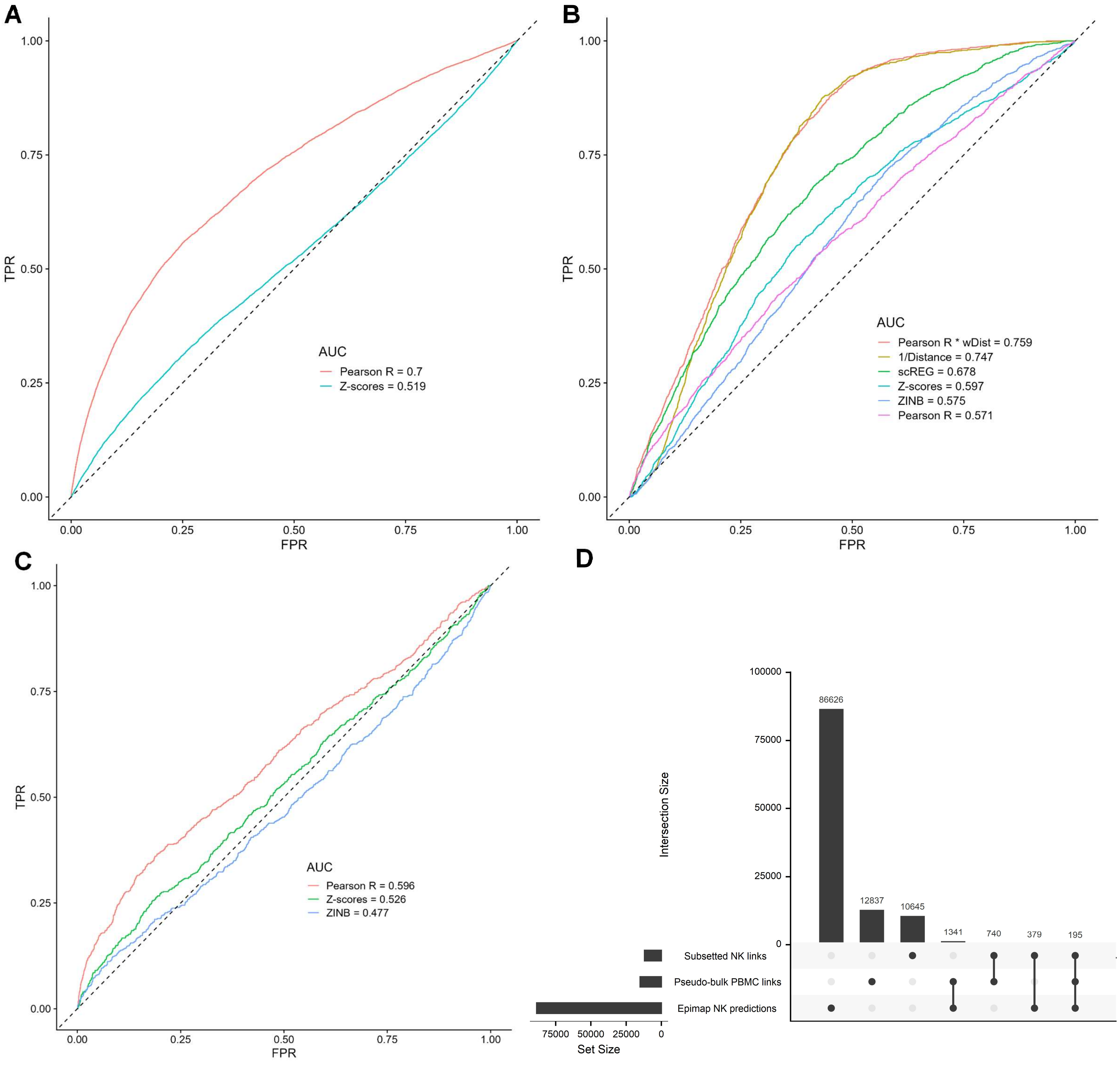

A
TPR
FPR
AUC
Pearson R = 0.7
Z-scores = 0.519
B
AUC
Pearson R * wDist = 0.759
1/Distance = 0.747
scREG = 0.678
Z-scores = 0.597
ZINB = 0.575
Pearson R = 0.571
C
AUC
Pearson R = 0.596
Z-scores = 0.526
ZINB = 0.477
D
Intersection Size
86626
12837
10645
1341
740
379
195
Subsetted NK links
Pseudo-bulk PBMC links
Epimap NK predictions
Set Size

**Supplementary Figure 9.** The Pearson R method more accurately validates Epimap-predicted links between cCRE and target genes in NK cells. (**A**) We used the Pearson R and Z-scores methods to detect links between ATACseq peaks and target genes (590,842 links with |Pearson R| >0.01) in the complete (i.e. pseudo-bulk) PBMC multiomic dataset. Then, we performed Receiving Operating Curves (ROC) analyses to compare the identified peak-gene links from the multiomic data with regulatory links in NK cells predicted by the Epimap Project. (**B**) As in **A**, but using a smaller set of links defined using a more stringent statistical threshold (15,113 links with |Pearson R| >0.1). All cell-types are used to identify links (pseudo-bulk), except for scREG which by design output link scores by cell-type (in this case, NK cells). (**C**) As in **B**, but limiting these ROC analyses to links between ATACseq peaks and target genes with |Pearson R| >0.1 that were found in the NK cells subset of the PBMC multiomic dataset. (**D**) Upset plot that shows the intersections of links identified between ATACseq peaks and target genes using either the full PBMC multiomic dataset (i.e. pseudo-bulk) or only the NK cells subset with cCRE-gene regulatory links in NK cells as predicted by the Epimap Project. ZINB; zero-inflated negative binomial, wDist; weighted distance ($e^{(-distance/200kb)}$), TPR, true positive rate; FPR, false positive rate.
